# Endomucin knockout leads to delayed retinal vascular development and reduced ocular pathological neovascularization

**DOI:** 10.1101/2024.07.16.603729

**Authors:** Zhengping Hu, Issahy Cano, Anton Lennikov, Melissa Wild, Urvi Gupta, Yin Shan Eric Ng, Patricia A. D’Amore

## Abstract

Endomucin (EMCN), an endothelial-specific glycocalyx component highly expressed in capillary and venous endothelium, plays a critical role in regulating VEGF receptor 2 (VEGFR2) endocytosis and downstream VEGF signaling. Using the first global EMCN knockout mouse model, we investigated the effects of EMCN deficiency on retinal vascularization during development and pathological angiogenesis. We found relatively high expression of EMCN in choroidal capillaries and retinal vasculature. Emcn^-/-^ mice exhibited delayed retinal vascularization at postnatal day 5, with fewer tip cells and reduced vessel density. Ultrastructural examination revealed disrupted and reduced fenestrations in choroidal capillary endothelium. In an oxygen-induced retinopathy model, while Emcn^-/-^ mice showed no significant difference in avascular area compared to Emcn^+/+^ mice at postnatal day 12, there was a significant reduction in neovascular tufts in Emcn^-/-^ mice at postnatal day 17. Similarly, in a laser-induced choroidal neovascularization model, Emcn^-/-^ mice showed a significant reduction in vascular leakage and lesion size. These findings suggest that EMCN plays a critical role in both vascular development and pathological neovascularization, highlighting its potential as a target for anti-angiogenic therapies.

## Introduction

Angiogenesis, the formation of new capillaries from existing blood vessels by branching^1^, is central to a variety of physiological and pathological processes, including embryonic development, wound healing, tumor growth and metastasis, and several ocular diseases such as wet age-related macular degeneration, proliferative diabetic retinopathy, and retinopathy of prematurity^2–7^. During angiogenesis, vessels are formed by budding and sprouting of endothelial cells (ECs), which play a crucial role in this precisely orchestrated process, including migration, proliferation, lumen formation, and final integration of nascent vessels with adjacent sprouts^8, 9^. Venous ECs exhibit distinctive characteristics, serving as the primary source of ECs during developmental and pathological angiogenesis^8, 10, 11^.

A variety of growth factors are involved in angiogenesis, and vascular endothelial growth factor A (VEGF) plays an essential role in regulating physiological and pathological angiogenesis, vascular permeability, and cell survival^12^. VEGF interacts with three tyrosine kinase receptors: VEGF receptor 1 (VEGFR1), VEGFR2, and VEGFR3. Among these receptors, VEGFR2 has a primary role in mediating the diverse biological effects of VEGF^13^. The binding of VEGF to VEGFR2 initiates a cascade of signaling events leading to tip cell selection and subsequent vascular branching^14^. VEGF binding to VEGR2 induces receptor dimerization, autophosphorylation, and internalization, resulting in increased tyrosine kinase activity and downstream signaling cascades, including the ERK1/2 and PI3k/Akt pathways^15, 16^.

The endothelial glycocalyx, a carbohydrate-rich layer lining the luminal surface of ECs, consists of glycoproteins, glycolipids, and proteoglycans. Recent research has highlighted the crucial role of the endothelial glycocalyx in regulating angiogenesis. Accumulating evidence suggests that the endothelial glycocalyx and its components play a pivotal regulatory role in angiogenesis^17–21^. Furthermore, specific glycocalyx elements have been found to interact with important angiogenic signaling pathways. Endomucin (EMCN), a type I transmembrane glycoprotein of the endothelial glycocalyx, is expressed on the surface of capillary and venous endothelium. EMCN has been widely used as an endothelial marker^22^ and its possible functional roles have only recently begun to be explored. Tissues with high vascularity, such as the kidney lung and the choroid, exhibit robust EMCN expression^23^. We previously reported a role for EMCN as an anti-adhesive molecule, preventing interactions between neutrophils and ECs^24^. In addition, accumulating evidence supports the involvement of EMCN in angiogenesis^25, 26^. The expression of EMCN by ECs is increased during proliferation and in response to stimulation with tumor-conditioned media^22^. We have previously shown that cystic embryoid bodies derived from VEGF-null murine embryonic stem cells exhibit reduced EMCN expression and impaired formation of vascular-like structures^25^. We demonstrated that EMCN knockdown in vitro inhibits VEGF-induced EC cell proliferation, migration, and tube formation^26^. We have further revealed an interaction between VEGFR2 and EMCN that is important in the regulation of VEGFR2 endocytosis^22, 27^.

In this study, the role of EMCN in vascular development and angiogenesis in the eye was characterized using an global EMCN knockout mouse model. Deletion of the EMCN gene in mice resulted in delayed retinal vascularization, reduced choriocapillaris fenestrae, decreased neovascularization area in the oxygen-induced retinopathy model, and smaller lesion size in the laser-induced choroidal neovascularization model. These findings support the function of EMCN in the regulation of angiogenesis in vivo, highlighting its potential role as a therapeutic target for angiogenesis-related diseases.

## Methods and materials

### Animals

Experimental procedures were conducted following the guidelines of and with the approval of the Institutional Animal Care and Use Committee at Schepens Eye Research Institute Massachusetts Eye and Ear (SERI-MEE). All protocols adhered to the ethical standards outlined by the Association for Research in Vision and Ophthalmology (ARVO) for the care and use of animals. The mice used in the study were raised under a 12-hour on/12-hour off-cyclic lighting schedule. Emcn-floxed mice were generated in conjunction with Cyagen Biosciences. A reporter gene (EGFP) was expressed upon successful Cre recombination and deletion of the EMCN gene. Global homozygous EMCN knockout (Emcn^-/-^) mice were obtained by breeding Emcn^flox/flox^ mice with ROSA26-Cre mice. After confirmation of successfully recombination of Cre, Emcn^+/-^, Cre was then back cross with Emcn^+/+^ to generate Emcn^+/-^ without Cre. Emcn^+/-^ male and female were used for breeding, and Emcn^+/+^, Emcn^+/-^ and Emcn^-/-^ littermates were used in this study. To confirm the deletion of EMCN in various tissues, including ear snip, retina, choroid, kidney, lung, thyroid, spleen, liver, and small intestine, were collected from Emcn^+/+^, Emcn^+/-^ and Emcn^-/-^ mice (8-to 16-wk-old), total RNA was extracted, and levels of Emcn mRNA assessed.

### Reagents and antibodies

Lysis Buffer (#9803S) and protease inhibitors (#5871S) were procured from Cell Signaling Technology. Additional reagents, including Tween-20 (#X251-07), phosphate-buffered saline (PBS, #D5652-10×1L), DL-dithiothreitol (DTT, #D9163-5G), and bovine serum albumin (BSA, #A6003), were sourced from Sigma-Aldrich. Protein standards used were Precision Plus Protein Dual Color Standards (#161-0374) from Bio-Rad.

BioTrace™ NT Nitrocellulose Transfer Membrane 30 cm x 3 m roll (#27376-991), DNA Gel Loading Dye (#R0611), and Fisherbrand™ RNase-Free Disposable Pellet Pestles (#12-141-364) were acquired from Fisher Scientific. For RNA stabilization, Invitrogen™ RNAlater™ Stabilization Solution (#AM7020) was used, obtained from Invitrogen. RNA extraction kits, including the RNase-Free DNase Set (#79254), RNeasy Plus Mini Kit (#74136), and QIAamp Fast DNA Tissue Kit (#51404), were supplied by Qiagen.

For immunoblotting, the membranes were probed with primary antibodies rat anti-mouse EMCN (1:1,000, Abcam, ab106100) and mouse anti-tubulin (1:2,000, Cell Signaling, #3873S). Tissue sections from frozen and paraffin-embedded eye cups were subjected to immunohistochemistry using the following primary antibodies: rat anti-mouse EMCN (1:200, Abcam, #ab106100), rabbit anti-EGFP (1:300, Sigma, #SAB4701015), and mouse anti-CD31 (1:200, Cell Signaling, #3528S).Secondary antibodies used for immunohistochemistry included donkey anti-rat Alexa Fluor 594 (1:300, Thermo Fisher, #A21209), goat anti-rabbit Alexa Fluor 488 (1:300, Thermo Fisher, #A11008), donkey anti-goat IgG Alexa Fluor 647 (1:300, Thermo Fisher, #A32849), and goat anti-mouse Alexa Fluor 647 (1:300, Thermo Fisher, #A21235).

### Analysis of mRNA expression

Freshly dissected retinas and RPE/choroid complexes were collected in RNAlater stabilization solution (Invitrogen, #AM7020). Total RNA was extracted using the RNeasy Mini Kit (Qiagen, #74136). Five hundred nanograms of purified RNA was reverse transcribed into cDNA using SuperScript IV reverse transcriptase (Invitrogen, #11756050). PCR amplification reactions, starting with 10-20 ng of cDNA, were performed on the LightCycler 480 II (Roche, Indianapolis, IN, USA) using FastStart Universal SYBR Green master mix (Roche, #A25743) and 0.4 μM primers. All samples were prepared with equal volumes of starting reagents. Relative mRNA levels were quantified using the Delta Delta Ct method, normalizing first to the average of the housekeeping genes Hprt1 and B2m, and then comparing the resulting Delta Ct values to those from wild-type controls. For qPCR analysis, all samples were prepared with the same amount of total RNA and equal volumes of reagents.

### Protein extraction and western blots

Freshly collected tissues were either preserved in Allprotect Tissue Reagent (#76405) or lysed using RIPA buffer (Cell Signaling, #9806S) supplemented with Protease Inhibitor Cocktail (1:100, Cell Signaling, #5871S), followed by homogenization on ice using a homogenizer (Fisher Scientific, #12-141-361). After centrifugation at 12,000g for 5 minutes in a benchtop centrifuge, supernatants were prepared to ensure equal protein concentrations, determined using the Pierce BCA assay kit (Thermo Scientific, #23227). Laemmli’s SDS Sample Buffer (Boston Bio Products, #BP-110R) was added to the samples, which were then boiled at 95°C for 5 minutes. Subsequently, the samples were separated by electrophoresis for 2 hours at 60-100 V using 4% to 20% precast gradient gels (Mini-PROTEAN TGX; Bio-Rad, #4561094) in 1X SDS-Tris-Glycine buffer (Bio-Rad, #1610772EDU). The gradient gels were transferred onto 0.45-μm nitrocellulose membranes (VWR, #27376-991) for 1 hour at 75 V using ice-cold 20% methanol in 1X Tris-glycine buffer (Bio-Rad, #1610771EDU). Following transfer, membranes were blocked for 1 hour with 3% BSA in PBS and then probed overnight at 4°C with primary antibodies targeting the proteins of interest. For detection, appropriate secondary antibodies, such as IRDye® 800CW goat anti-rabbit (Licor, #925-32211) or IRDye® 680RD goat anti-mouse (Licor, #925-68070), were utilized. Final image development was performed using the LI-COR Odyssey fluorescence system (LI-COR). Band intensities were quantified using ImageJ software.

### Immunohistochemistry

Eye cups were dissected from freshly collected eyes and fixed in 4% PFA overnight at 4°C. Paraffin sections were prepared by the Morphology Core at SERI-MEE and stained with H&E. Frozen sections of eye cups were used for immunohistochemistry. Slides were initially treated with a 0.3% H2O2 solution at room temperature for 10 minutes to block endogenous peroxidase activity, followed by washing with PBS and distilled water. Slides were then permeabilized with 0.5% Triton X-100 in PBS for 5 minutes and blocked with 5% goat or donkey serum in PBS for 1 hour at room temperature.

Primary antibodies, prepared in blocking buffer, were applied to the slides and incubated overnight at 4°C in a humidified chamber. The sections were subsequently washed with PBS and incubated with secondary antibodies in blocking buffer for 2 hours at room temperature. Slides were mounted with Prolong Gold Antifade Reagent containing DAPI (Invitrogen, #P36935) and visualized using a Zeiss Axioscope 7 microscope (Oberkochen, Germany).

### Development retinal vascularization

Pups at postnatal day 5 (P5) from Emcn^+/-^ and Emcn^+/-^ breeders were euthanized, and the eyes were collected for fixation in 4% paraformaldehyde (PFA) at 4°C overnight. Ear snips were also obtained from the pups for genotyping. The retinas were then flat-mounted and stained with isolectin-B4. Imaging of the retinal vasculature was performed using a Zeiss Axioscope 7 microscope (Oberkochen, Germany) with automatic stitching. The angiogenic fronts of the P5 retina flat mounts were further examined using Leica TCS SP8 Confocal Microscopy (Leica Microsystems, Wetzlar, Germany). The area of retinal vessels over the entire retina area was quantified using Image J from retinal flat mount images. Tip cells and vessel branches/segments were quantified using the angiogenesis plugin on Image J. Genotypes of the pups were determined after image quantification.

### Oxygen-induced retinopathy (OIR)

We conducted the OIR model as previously described^28^. Up to four cages, each housing a nursing mother and her P7 litter, were placed in a chamber set at 75% oxygen concentration for five days. Subsequently, at P12, the animals were returned to room air conditions for another five days. Oxygen levels within the chamber were monitored daily using an independent oxygen sensor (Advanced Instruments, GPR-20F). To mitigate the risk of acute lung toxicity, the dam of the experimental animals was rotated daily with a dam kept under normal oxygen conditions for the duration of five days. Pups were euthanized at both P12 and P17, and their retinas were carefully dissected, flat-mounted, and subjected to immunostaining. The avascular areas at P12 and the neovascular areas at P17 were quantified using Adobe Photoshop 2022 software, as described previously^29^. Additionally, retinas harvested at P17 were processed for RNA extraction to analyze mRNA expression levels of angiogenesis-related genes.

### Laser induced choroidal neovascularization (CNV)

Six-to 8-wk–old Emcn^+/+^, Emcn^+/-,^ and Emcn^-/-^ mice were included in CNV model. Mice were anesthetized by intraperitoneal injection of a ketamine/xylazine mixture (100/50 mg/kg). Laser photocoagulation was performed under general anesthesia with a 532-nm laser attached to the Micron III image-guide system (Phoenix Technology Group, Pleasanton, CA) using 150 mW power, 50-ms duration, and a spot size of 50 μm. Four laser spots were placed around the optic nerve. Disruption of Bruch’s membrane was confirmed by the appearance of a cavitation bubble. Seven days after laser induction, fluorescein angiography was conducted following injection of 0.1 mL of 2% sodium fluorescein, optical coherence tomography (OCT) was performed, and serial photographs were captured. Light source intensity and gain were standardized and maintained for imaging in all experiments. Animals were euthanized, and eyes were collected on day eight post-laser for choroidal flat mount and immunostaining. Vessel leakage was measured by lesion intensity late time point (5 min) - lesion intensity early time point (30 sec). Lesion size was quantified using FA images. Lesion area was quantified using OCT images, and lesion size was quantified using IB4 stained choroidal flat-mount. All quantification was conducted in a masked fashion using Image J.

### Retina and choroidal flatmounts and immunostaining

Eyes were enucleated and fixed in 4% PFA overnight at 4°C. The cornea and lens were removed, and the retinas or RPE/choroid complexes were flatmounted by cutting the tissues into 4-8 petals. The tissues were blocked with 5% bovine serum albumin, 0.1% Triton X-100, and 3% donkey serum in PBlec buffer overnight at 4°C, incubated with 594 conjugated-IB4 (1:200, Thermo Fisher Scientific), or mouse anti-VEGF (1:100; #sc-7269, Santa Cruz) and the corresponding secondary antibody, and then mounted on slides with ProLong gold antifade mounting media(Life Technologies, #P36935). Images were acquired with Zeiss Axioscope 7 microscope (Oberkochen, Germany) or Leica TCS SP8 Confocal Microscopy (Leica Microsystems, Wetzlar, Germany).

### Transmission electron microscopy

Mice were perfused with PBS, followed by half Karnovsky’s fixative (2% formaldehyde + 2.5% glutaraldehyde, in 0.1 mol/L sodium cacodylate buffer, pH 7.4). Eyes were collected and immersed in half Karnovsky’s fixative for 24 hr at 4°C. After fixation, samples were processed by Morphology Core at SERI-MEE for TEM imaging. In brief, the eye cups were trimmed into 1-mm thick segments and rinsed with 0.1 mol/L sodium cacodylate buffer. They were then post-fixed and stained en bloc with 2% gadolinium triacetate in 0.05 mol/L sodium maleate buffer (pH 6) for 30 minutes. After dehydration and polymerized using a 60°C oven, semithin sections (1 μm thickness) were cut and stained with 1% toluidine blue in a 1% sodium tetraborate aqueous solution for examination by light microscopy. Ultrathin sections were obtained and stained with aqueous 2.5% gadolinium triacetate and modified Sato lead citrate and were imaged using an FEI Tecnai G2 Spirit transmission electron microscope (FEI, Hillsboro, OR) at 80 kV interfaced with an AMT XR41 digital charge-coupled device camera (Advanced Microscopy Techniques, Woburn, MA) for digital image acquisition at 2000 × 2000 pixels at 16-bit resolution. The length of fenestrated endothelium on choriocapillaris and the number of fenestrae were quantified using Image J in a masked manner.

### Statistical analysis

All values are expressed as mean ± SEM. Statistical analysis was performed using an unpaired Student t-test, one-way-ANOVA (GraphPad Prism 9). A P value <0.05 was considered statistically significant. Each experimental condition was conducted at least in triplicate, and all experiments were independently repeated at least three times.

## Result

### Choriocapillaris endothelium express a high level of EMCN

The levels of Emcn mRNA in various tissues, including the retina, choroid/RPE, liver, small intestine, kidney, spleen, thyroid, and lung, were examined (Figure 1A). Emcn mRNA levels were first normalized to housekeeping genes (average of B2m, Hprt1) and then compared to Emcn levels in the spleen. Among all the tissues examined, the choroid/RPE had the highest levels of Emcn mRNA (retina: 8.9 ± 1.7; choroid: 235.0 ± 2.14; liver: 4.3 ± 1.3; small intestine:1.7 ± 0.31; kidney:143.6 ± 16.7; thyroid: 5.5 ± 15.8; lung: 50.2± 20.6). To confirm the deletion of Emcn in Emcn^-/-^ mice, choroid/RPE complexes were collected from Emcn^+/+^, Emcn^+/-^ and Emcn^-/-^ and EMCN mRNA (Figure 1B) and protein (Figure 1C) measured. EMCN levels were significantly reduced by about 50% in Emcn^+/-^ and were undetectable in Emcn^-/-^ compared to Emcn^+/+^. The localization of EMCN was examined using immunohistochemistry on frozen sections of eye cups from Emcn^+/+^, Emcn^+/-^, and Emcn^-/-^ mice (Figure 1D). EMCN was expressed by retinal vasculature, highly expressed by choroidal capillaries and co-localized with endothelial marker CD31. EGFP, a reporter of EMCN deletion, was detected in Emcn^+/-^ and Emcn^-/-^ and co-localized with CD31. EMCN was not detected in Emcn^-/-^ eyecups.

**Figure 1.**
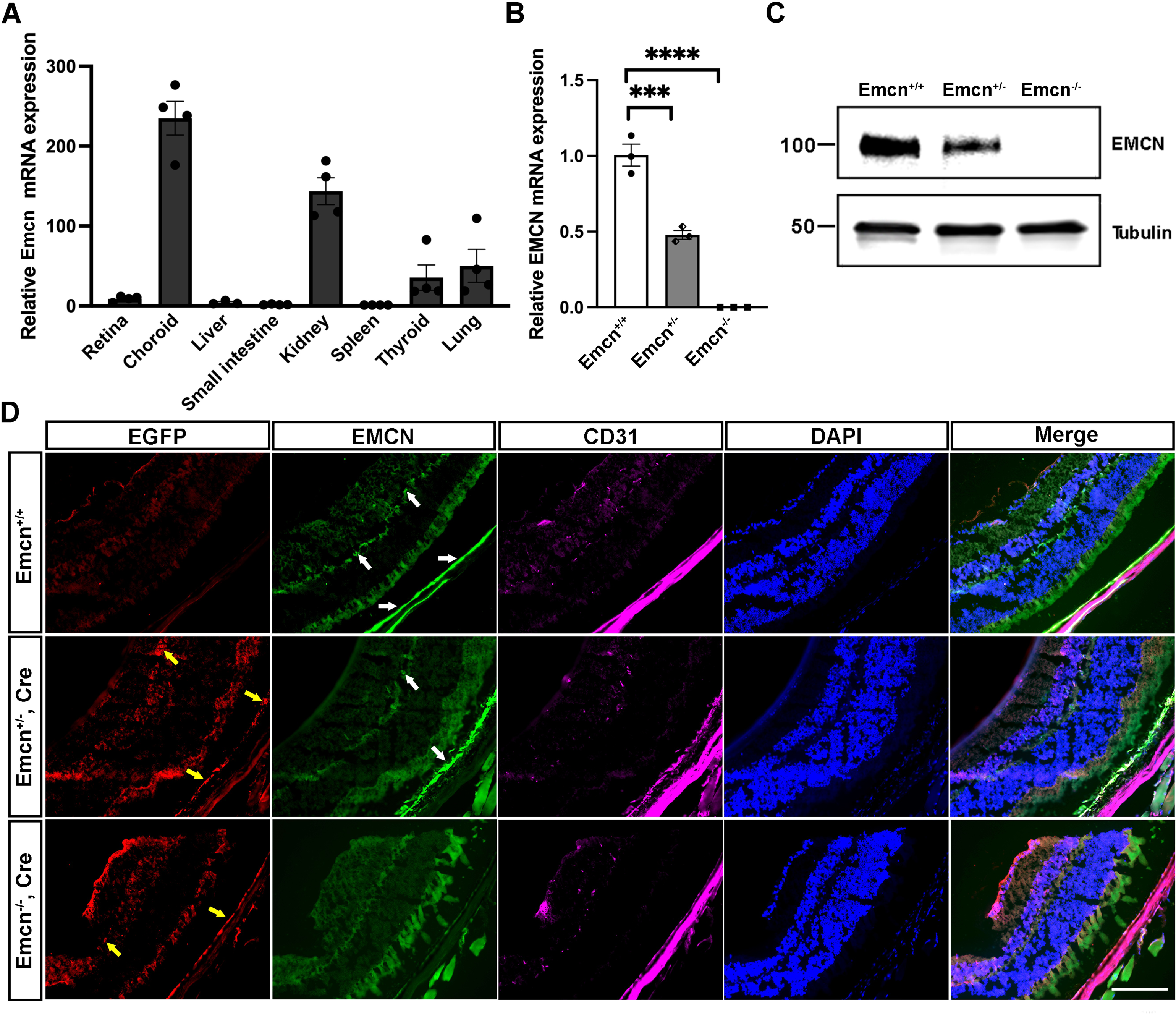
Choroid expresses high levels of EMCN. **(A)** Retina, choroid/RPE complex, kidney, lung, thyroid gland, spleen, small intestine, and liver tissues were collected from wild-type mice, and mRNA was extracted and examined for EMCN expression. n=4. **(B)** mRNA was collected from freshly harvested choroid/RPE complexes from Emcn^+/+^, Emcn^+/-^, and Emcn^-/-^ mice and examined for EMCN expression. ***, p<0.0001, ****, p<0.0001, n=3, One-way ANOVA. **(C)** Proteins were extracted from choroid/RPE complexes from Emcn^+/+^, Emcn^+/-^, and Emcn^-/-^ mice and quantified using the BCA protein assay kit. Thirty 30 μg of each sample were loaded onto an SDS-PAGE gel. Levels of EMCN and tubulin were examined by western blot. **(D)** Expression and localization of EMCN, eGFP, CD31, and nuclear DAPI were examined in sections of retina/choroid by immunohistochemistry in Emcn^+/+^, Emcn^+/-^, and Emcn^-/-^ mice. Yellow arrows indicate EGFP expression and white arrows indicate EMCN expression. Scale bar = 100 µm.

### EMCN deletion leads to delayed retinal vascular development

To investigate the role of EMCN in the developing retinal vasculature, eyes were collected from Emcn^+/+^, Emcn^+/-,^ and Emcn^-/-^ littermates at postnatal day 5 (P5) for retinal flat-mounts. Isolectin B4(IB4) stained the retinal vasculatures at P5 (Figure 2A). The masked quantification of the ratio of vessel area to the retinal area revealed a significant decrease in retinal vascularization in both Emcn^+/-^ and Emcn^-/-^ compared to the Emcn^+/+^ (0.1770 ± 0.01 and 0.145 ± 0.01 vs. 0.201 ± 0.01, p<0.001 and p<0.00001, Welch’s t-test) (Figure 2B). Higher magnification images of the angiogenic front of P5 retinal vessels examined by confocal shown in Figure 2C show a lower density of retinal vessels in Emcn^-/-^ compared to Emcn^+/+^ as well as a reduction in the number of endothelial tip cells (13.00 ± 1.3 vs. 16.00 ± 1.3, p>0.05; 11.50 ± 1.2 vs. 16.00 ±1.3, p<0.05. One-way ANOVA) (Figure 2D). Image J quantification of the vessel branches (Figure 2E) and segments (Figure 2F) of the P5 angiogenic front indicated significant reductions in both Emcn^+/-^ and Emcn^-/-^ mice compared to Emcn^+/+^ (branches: 32.0 ± 1.6 vs. 40.2 ± 1.3 p<0.005; 29.8 ± 1.3 vs. 40.2 ± 1.3, p<0.0005; segments: 365.5 ± 37.32vs. 456.3 ± 8.8, p<0.05, 330.3 ± 16.8 vs. 456.3± 8.8, p<0.005, One-way ANOVA).

**Figure 2.**
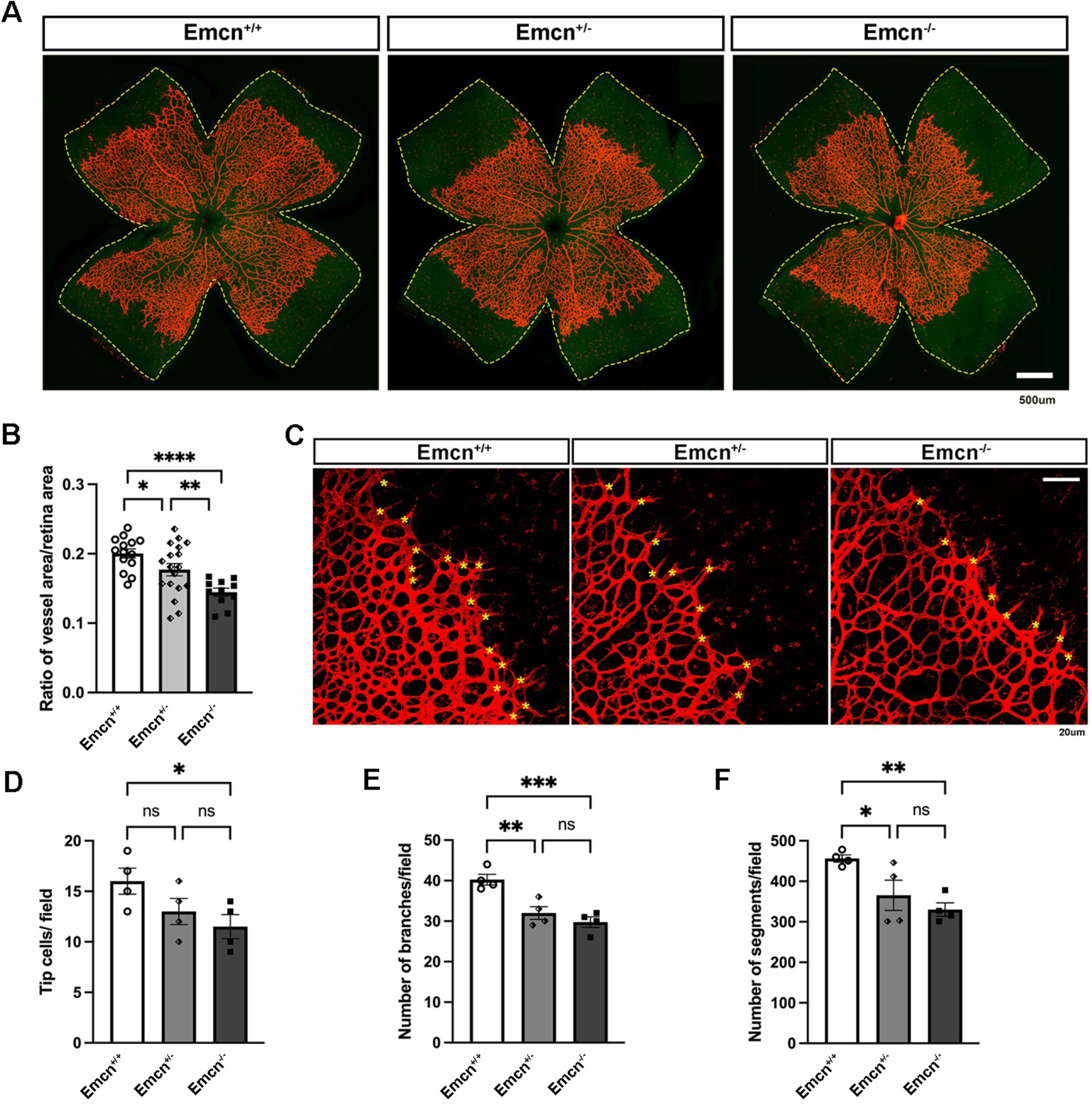
Delayed retinal vascular development at P5 in Emcn^-/-^mice. **(A)** Eyes from P5 mice and adult mice were collected for retinal flat mounts. Retinal vasculature (superficial layer) was stained with isolectin-B4 (IB4) and imaged with a ZEISS Axioscope. Scale bar = 500 µm. **(B)** Quantification of the retinal vascular area and total retina area were conducted in a masked fashion using ImageJ. * p<0.01, ** p<0.001, **** p<0.00001. n=14,18,11, Welch’s t-test. **(C)** Representative images of the angiogenic fronts from P5 IB4-stained retinal flat mount were obtained using confocal microscopy (SP8). Yellow stars mark the tip cells. Scale bar = 20 µm. **(D-F)** Masked quantification of tip cells per field, the number of branches and segments per field of the P5 angiogenic front images from Emcn^+/+^, Emcn^+/-^, and Emcn^-/-^ mice. *, p<0.05,**, p<0.001,***, p<0.0001, n=4, One-way ANOVA.

### The choriocapillaris of Emcn^-/-^ mice display disrupted fenestrae

To examine the morphology of Emcn KO eyes, we performed H&E staining on paraffin sections of Emcn^+/+^, Emcn^+/-^, and Emcn^-/-^ mice (Figure 3A). No significant differences were observed in the thickness of different retinal layers. We then conducted ultrastructural analysis of choroidal capillaries in Emcn^+/+^, Emcn^+/-^, and Emcn^-/-^ mice.

**Figure 3.**
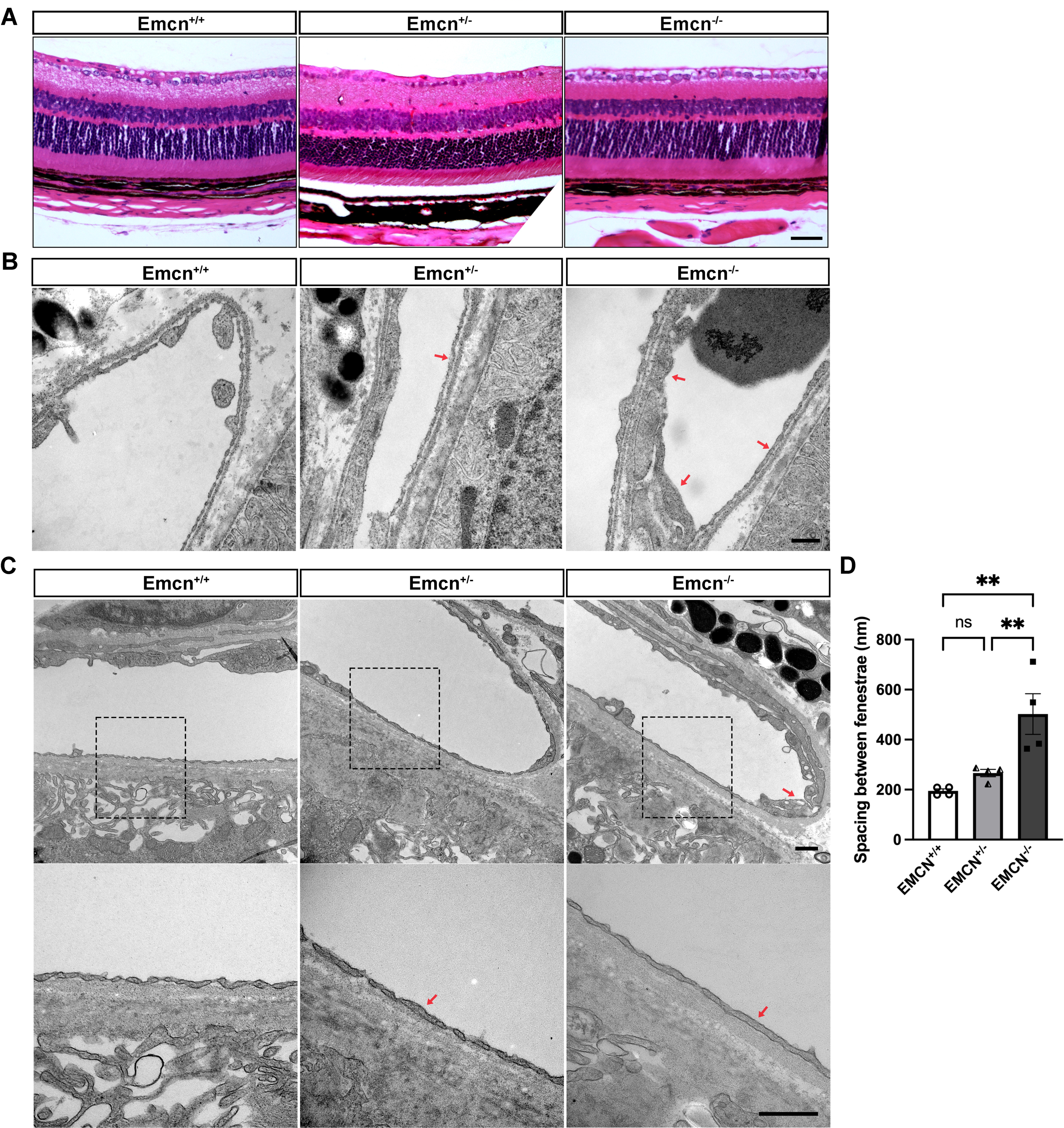
Reduction of fenestrae in choroidal capillaries in Emcn^-/-^ mice. **(A)** H-E staining of paraffin sections of eye cups collected from adult Emcn^+/+^, Emcn^+/-,^ and Emcn^-/-^ mice. Representative images of TEM from eyes collected from perfuse-fixed **(B)** adult (4 to 6-month-old) or **(C)** aged (20 to 24-month-old) Emcn^+/+^, Emcn^+/-^ and Emcn^-/-^ mice. Red arrows indicate areas of choroidal capillaries that lack fenestrae. N = 4. Scale bar = 500 nm **(D)** Masked quantification of the distance between fenestrae in Emcn^+/+^, Emcn^+/-^ and Emcn^-/-^ mice using ImageJ. *, p<0.05, n=4, One-way ANOVA.

TEM images from the adult mice (4 to 6-month-old) revealed disorganized fenestrations in the choriocapillaris and regional thickening of endothelial cells in EMCN^-/-^ mice compared to wild-type (Figure 3B). TEM images from the eyes of the aged mice (20 to 24-month-old) indicated a further disruption of choriocapillaris fenestrae in both Emcn^+/-^ and Emcn^-/-^ mice (Figure 3C). Masked quantification of the average spacing between fenestrae on Emcn^+/+^, Emcn^+/-^, and Emcn^-/-^ mice showed significant increase in spacing Emcn^-/-^ compared to both Emcn^+/+^ and Emcn^+/-^ mice (502.1 ± 81.2 nm vs. 194.7 ± 8.2 nm, p<0.001; 502.1 ± 81.2 nm vs. 266.3 ± 14.6 nm, p<0.001, One-way ANOVA).

### EMCN knockout results in reduced neovascular area in the OIR model

Next, we investigated the involvement of EMCN in the neovascularization of the OIR model. First, the avascular area of the retina flat mounts from Emcn^+/+^, Emcn^+/-^, and Emcn^-/-^ mice were examined following their return to room air at P12 (Figure 4A). Masked quantification of the percentage of avascular area in the retina indicated no significant difference among Emcn^+/+^, Emcn^+/-^, and Emcn^-/-^ mice (23.2 ± 0.86%, 24.10 ± 0.6%, and 24.57 ± 0.60%, p>0.05, One-way ANOVA) (Figure 4B). Pathological neovascularization at P17 was also examined (Figure 4C). Masked quantification of the percentage of the total neovascular area showed a significant reduction in Emcn^+/-^ and Emcn^-/-^ mice compared to Emcn^+/+^ mice (7.74 ± 0.7% vs. 12.0 ± 1.4%, p<0.05; 9.0 ± 2.9% vs. 12.0 ± 1.4%, p<0.05, One-way ANOVA) (Figure 4D). We further examined the neovascular tufts as well as the presence of microglia/macrophage on P17 retinal flat mounts using immunohistochemistry (Figure 5A). The majority of microglia/macrophage cells were localized adjacent to the areas of neovascularization. The staining of retinal vessels using IB4 allowed for better visualization of the neovascularization in each retinal flat mount quadrant. qPCR analysis revealed a significant reduction in the relative expression of VEGF (0.53 ± 0.1 vs. 1.00 ± 0.2, p<0.05; 0.5 ± 0.1 vs. 1.06 ± 0.1, p<0.05, One-way ANOVA) (Figure 5B) and VEGFR2 (0.54 ± 0.1 vs. 1.00 ± 0.1, p<0.05; 0.5 ± 0.05 vs. 1.4 ± 0.1, p<0.05, One-way ANOVA)(Figure 5C) in Emcn^-/-^ compared to Emcn^+/+^ and Emcn^+/-^ at mRNA level. No significant difference in was detected in the levels of HIF-1a mRNA among the Emcn^+/+^, Emcn^+/-^, and Emcn^-/-^ mice (1.00 ±0.2, 1.22 ± 0.2, 0.87 ± 0.1, p>0.05, One-way ANOVA) (Figure 5D).

**Figure 4.**
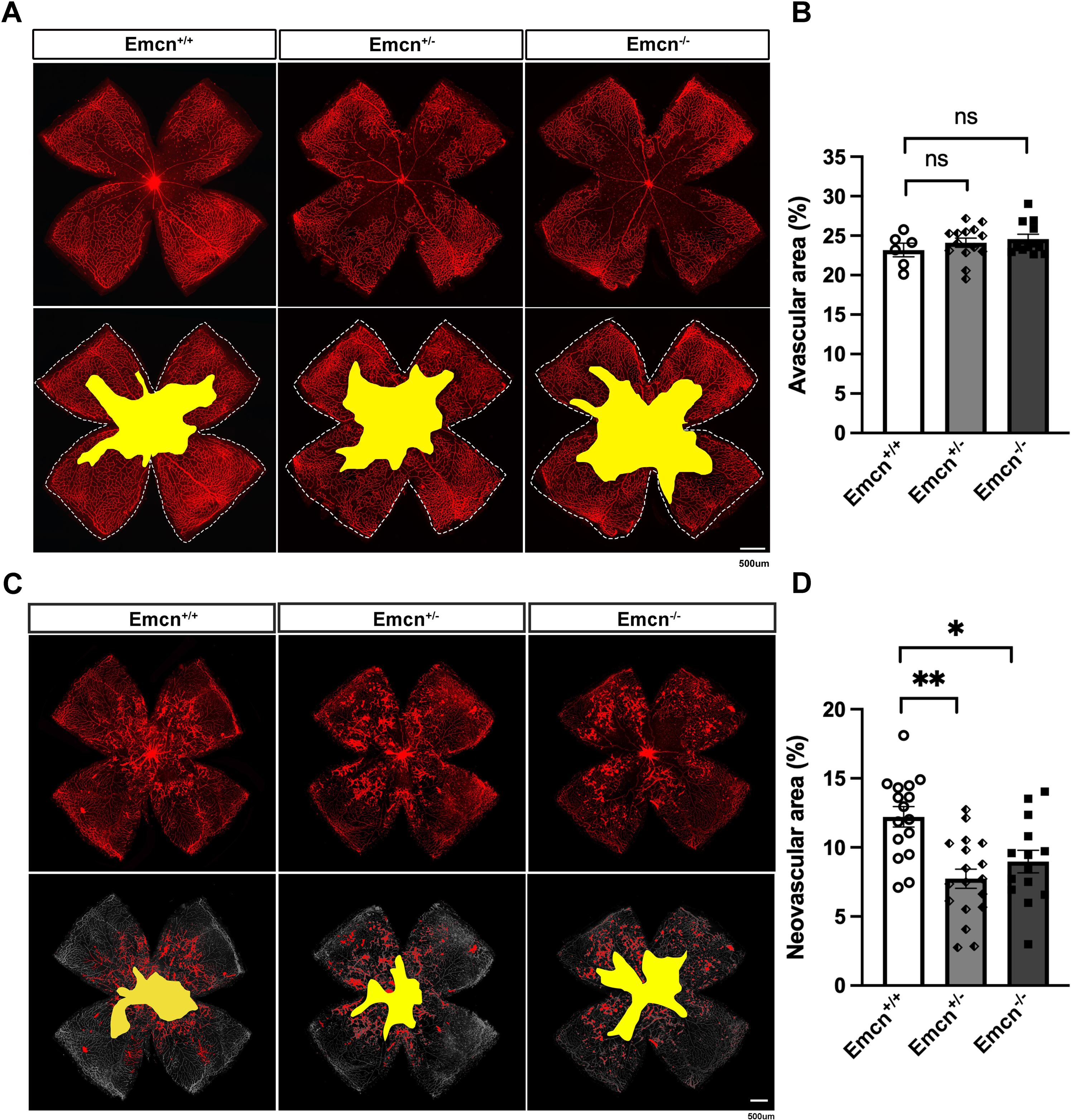
Reduced neovascular area in Emcn^-/-^ mice in the OIR model. **(A)** Representative images of the retinal vasculature at P12 stained with IB4. The avascular area is highlighted in yellow. Scale bar = 500 µm. **(B)** Avascular area of retinas of Emcn^+/+^, Emcn^+/-^, and Emcn^-/-^ mice at P12 were quantified in a masked fashion using Photoshop CC. n =6,14,13, ns, p>0.05. Welch’s t-test. **(C)** Representative images of retinal vasculature at P17 stained with IB4. The avascular area is highlighted in yellow, and the neovascular area is marked in red. Scare bar = 500 µm. **(D)** Neovascular areas at P17 from Emcn^+/+^, Emcn^+/-^, and Emcn^-/-^ mice were quantified using Photoshop CC in a masked fashion. Data = mean ± SEM, * p<0.05, ** p<0.001. n=12,17,14. Welch’s t-test.

**Figure 5.**
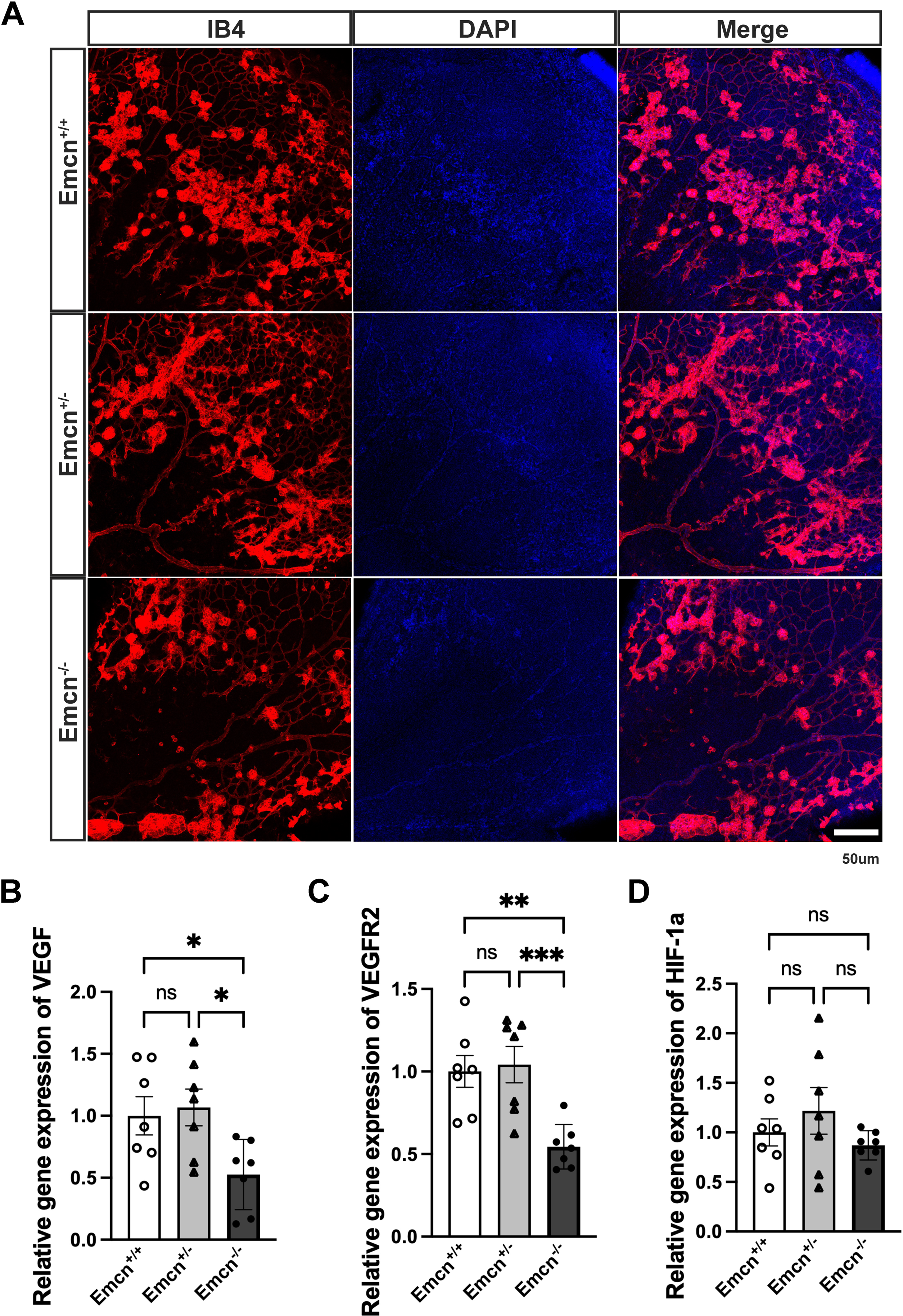
Reduced expression of VEGF and VEGFR2 expression in Emcn^-/-^ in the OIR model. **(A)** Representative confocal images of the neovascular area of P17 retinas from Emcn^+/+^, Emcn^+/-^, and Emcn^-/-^ mice stained with IB4 showing the neovascular tufts. **(B-D)** RNA was isolated from the retinas of P17 Emcn^+/+^ mice, and Vegf, Vegfr2, and Hif-1α gene expression was quantified by qPCR, normalized to housekeeping genes B2m and Hprt1. * p<0.01, ** p<0.001, *** p<0.0001. n=7, one-way ANOVA.

### Emcn^-/-^ mice develop smaller lesions in the laser-induced model of CNV

We then investigated the role of EMCN in the neovascularization in the laser-induced model of CNV^30, 31^. Figure 6A shows fundus images seven days after laser photocoagulation, along with fundus fluorescein angiography at both early and late time points, for Emcn^+/+^, Emcn^+/-^, and Emcn^-/-^ mice. Masked quantification of lesions leakage on FA revealed that Emcn^-/-^ mice had significantly reduced leakage compared to Emcn^+/+^ mice (22.2 x 10^4^ ± 2.2 x 10^4^ pixels vs. 41.06 x 10^4^ ± 3.6 x 10^4^ pixels, p<0.00001, 22.2 x 10^4^ ± 2.2 x 10^4^ pixels vs. 47.02 x 10^4^ ± 3.9 x 10^4^, p<0.0001, One-way ANOVA) (Figure 6B). The average lesion size quantified using the FA images at the late time point indicated smaller lesions in Emcn^-/-^ mice compared to both Emcn^+/+^ and Emcn^+/-^ mice (34.33 x 10^4^ ± 2.8 x 10^4^ pixels vs. 55.38 x 10^4^ ± 5.2 x 10^4^ pixels, p<0.001, 34.33 x 10^4^ ± 2.8 x 10^4^ pixels vs. 64.90 x 10^4^± 5.8 x10^4^, P<0.0001, One-way ANOVA) (Figure 6C). We performed the OCT on the Emcn^+/+^, Emcn^+/-^, and Emcn^-/-^ eye to visualize choroidal fibro-vascular tissue in a cross-sectional view (Figure 7A). The average area (four lesions in one eye) showed a reduction in Emcn^-/-^ mice compared to both Emcn^+/+^ and Emcn^+/-^ mice (2.68 x 10^4^ ± 0.1 x 10^4^ μm^2^ vs. 3.54 x 10^4^ ± 0.1 x 10^4^ μm, p<0.00001; 2.68 x 10 ± 0.1 x 10 μm vs. 3.24 x 10 ± 0.1 x 10 μm, p<0.001, One-way ANOVA) (Figure 7B). Masked quantification (Figure 7C) of the IB4 stained vessel area corroborated the OCT findings that Emcn^-/-^ mice had smaller lesions compared to Emcn^+/+^ and Emcn^+/-^ mice (2.2 x 10^4^ ± 0.2 x 10^4^ μm^2^ vs. 3.2 x 10^4^ ± 0.2 x 10^4^ μm^2^, p<0.001; 2.2 x 10^4^ ± 0.2 x 10^4^ μm^2^ vs. 3.3 x 10^4^ ±1.2 x 10^4^ μm^2^, p<0. 001. One-way ANOVA). Immunohistochemistry was further performed to visualize the vessels and VEGF expression in the CNV lesions on RPE/choroid flat-mounts (Figure 7D), which showed a significant reduction in lesions size in Emcn^-/-^, and Emcn^+/-^ compared to Emcn^+/+^.

**Figure 6.**
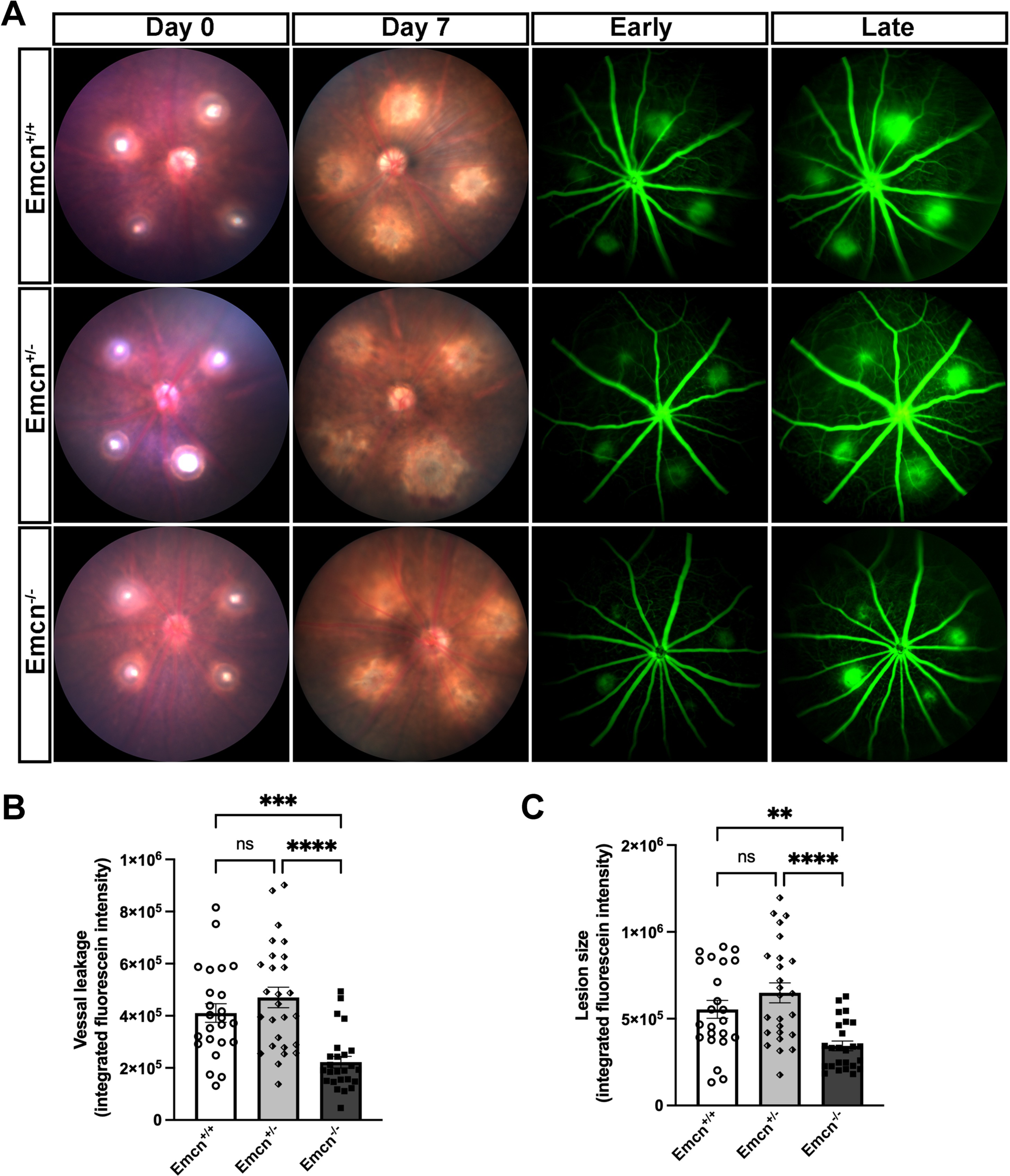
EMCN deletion leads to reduced lesion leakage in the laser-induced model of CNV. **(A)** Representative fundus images and FA from Emcn^+/+^, Emcn^+/-^, and Emcn^-/-^ mice at day seven after laser-induction of CNV. **(B)** The intensity of fluorescence leakage between early and late timepoint post sodium fluorescein injection in Emcn^+/+^, Emcn^+/-^, and Emcn^-/-^ mice were quantified in a masked fashion using ImageJ. n=24, 27,25, ***,p<0.0001,**** p<0.00001. Welch’s t-test (C) Masked quantification of lesion size in FA from the late time point on day seven in Emcn^+/+^, Emcn^+/-^, and Emcn^-/-^ mice. n=24, 27, 25, **, p<0.001, **** p<0.00001, Welch’s t-test.

**Figure 7.**
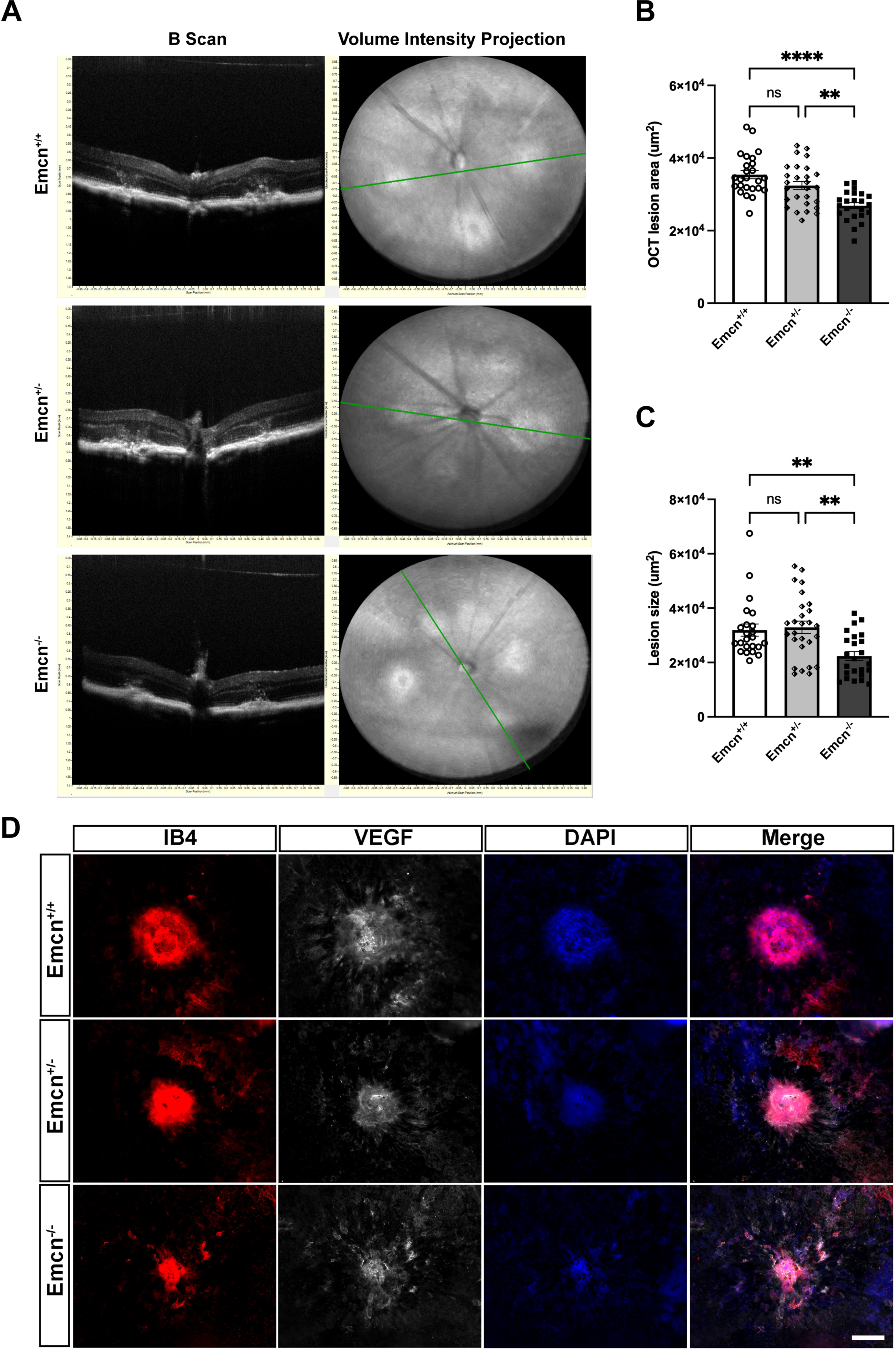
Emcn^-/-^ mice developed smaller CNV lesions. **(A)** Representative OCT images seven days following induction of laser-induced CNV. **(B)** Masked quantifications of lesion size from OCT imaging. Data = mean ± SEM, n=25, 27, 23, **** p<0.00001, Welch’s t-test **(C)** Quantification of lesion size from choroid/RPE flat mounts in Emcn^-/-^, Emcn^+/-^ compared to Emcn^+/+^. n=24, 27, 25, ** p<0.001, Welch’s t-test **(D)** Choroid/RPE flat mount from day eight following induction of CNV from Emcn^-/-^ mice, Emcn^+/-^ and Emcn^+/+^ control littermates (6 to 8-weeks-old) were stained for expression of IB4, and VEGF then imaged to visualize lesion size and VEGF expression. Scale bar = 50 µm.

## Discussion

Since its discovery in 1989^32^, VEGF, a signaling protein whose expression is known to be elevated in hypoxic environments^33^, has played a key role in the process of angiogenesis under developmental, physiological, and pathological conditions ^34, 35^. It exerts multiple effects on ECs, the key players in neovascularization^36^. VEGF expression in the retinal pigment epithelium is crucial for the development and maintenance of the choriocapillaris and overall visual function, as its absence leads to significant vascular and visual impairments^37^. EMCN, an endothelial specific glycoprotein, has been previously shown by our group to inhibits VEGF-induced ECs migration, growth, and morphogenesis by modulating VEGFR2 signaling^26^, and VEGFR2 endocytosis in vitro^27^. In this study, we advanced our understanding of the involvement of EMCN in vascular development and angiogenesis by using a EMCN knockout mouse model in the ocular setting. Our findings shed light on the potential of EMCN as a promising therapeutic target for anti-angiogenesis.

EMCN was originally identified as an endothelial marker^22^, and we together with other groups have confirmed that it is selectively expressed in venous and capillaries endothelium^23, 26, 38^. As we explored the expression of EMCN in various tissues, we observed a significantly high levels of EMCN in both the choroid and kidney. Notably, they shared the presence of fenestrations with diaphragm^39–42^. The development and maintenance of the fenestration dependent upon VEGF^43^. We have previously shown that EMCN expression is regulated by VEGF. Using gene expression microarray analysis, we observed that EMCN mRNA expression was about 3-fold higher in wild-type cystic embryonic bodies compared to VEGF-null cystic embryonic bodies, resulting in impaired vascular morphogenesis^25^. On the other hand, EMCN is closely involved in VEGF-induced signaling and downstream EC angiogenic functions. EMCN knockdown in vitro inhibits VEGF-induced VEGFR2 clathrin-mediated endocytosis^27, 44^, and VEGF-induced ECs migration, tube formation and proliferation^26^. The importance of EMCN extends to in vivo models as well, where its knockout has been demonstrated to disrupt vascularization in the developing mouse retina^26^. In the context of tip cell formation, which is essential for initiating new blood vessel sprouts, EMCN may influence the selection and behavior of these specialized endothelial cells as indicated by previous study that noted that Emcn+ bone marrow ECs expressed markers associated with endothelial tip cells and crucial for sprouting during vascular morphogenesis, such as Apln, Cxcr4, Kdr, Dll4, Angpt2, Sox17^45^. This is consistent with what we observed during the retinal vasculature development.

In our EMCN knockout mice, while no apparent retinal structural defects were observed through light microscopy in EMCN knockout mice, TEM analysis revealed notable endothelial thickening and loss of fenestration in choroidal capillaries compared to wild-type mice. This compelling data further confirms the critical role of EMCN in VEGF signaling and its involvement in maintaining the fenestrated phenotype in choroidal capillaries. Consistent with our observations in EMCN knockout mice, studies using VEGF-A blockade with VEGF-trap have shown similar changes in the choriocapillaris.

Specifically, regional endothelial thickening associated with loss of fenestration was observed^46^. These findings further support the importance of VEGF in maintaining the fenestrated endothelial phenotype and highlight the interplay between VEGF and EMCN in ECs fenestration. In adulthood, while EMCN is predominantly expressed on endothelial cells in vein, venous and capillary^47^, it continues to play a part in maintaining vascular homeostasis where it appears to be involved in regulating vascular permeability, leukocyte trafficking, and endothelial cell-cell interactions^48^.

Overexpressing EMCN in diabetic rats restores the endothelial glycocalyx, reduces inflammation and leukocyte adhesion, stabilizes the blood-retinal barrier, and inhibits vascular leakage^49^.

VEGF and its primary receptor VEGFR2 are central to the pathogenesis of neovascularization in both OIR and laser-induced CNV models. In OIR, the initial hyperoxic phase (75% O_2_, P7-P12) leads to downregulation of VEGF expression via oxygen-dependent degradation of hypoxia-inducible factor 1α (HIF-1α), resulting in endothelial cell apoptosis and vessel regression^50^. Upon return to room air (P12-P17), the relative hypoxia stabilizes HIF-1α, triggering a surge in VEGF expression^51^. This VEGF increase activates VEGFR2 on retinal endothelial cells, initiating several signaling cascades: the PLCγ/PKC/MAPK pathway promotes cell proliferation^52^, Src and FAK pathways enhance cell migration^53^, and the PI3K/Akt pathway supports cell survival^15^. Additionally, VEGFR2 signaling increases vascular permeability through eNOS activation and tight junction protein redistribution^54^. Our observations in the OIR model align with previous findings in the field. Specifically, we noted no significant difference in the avascular area at P12, but a significant reduction in neovascular tufts at P17. At mRNA level, we found no difference at HIF-1α level, but reduced VEGF and VEGFR2. This result suggests that the during the room air phase of the model, when VEGF levels typically surge, loss of EMCN exerts its effects on retinal endothelium. In the absence of EMCN, the internalization of VEGF receptors and subsequent downstream signaling in the endothelium is impaired^27^. Consequently, this leads to a reduction in pathological neovascularization, despite the elevated VEGF levels.

In the CNV model, laser injury induces an inflammatory response and local hypoxia, prompting VEGF production by RPE, inflammatory cells, and choroidal endothelial cells^55^. VEGF levels peak 3-7 days post-laser, driving CNV formation through VEGFR2-mediated endothelial cell activation, proliferation, and migration^56^. VEGFR2 signaling in CNV also increases vascular permeability^57^ and promotes endothelial cell survival via PI3K/Akt pathway activation^58^. Furthermore, VEGFR2 activation recruits bone marrow-derived cells, including endothelial progenitor cells, contributing to CNV progression^59^. Our findings in the laser-induced CNV model provide further compelling evidence for the critical role of EMCN in VEGF signaling. We observed a reduction in CNV lesion size as demonstrated by FA leakage, smaller lesions on OCT staining of the affected areas.

The pathology in both processes leads to the formation of disorganized, leaky vessels characteristic of pathological ocular neovascularization. While the two models differ in tissue origin, hypoxia dynamics, temporal patterns of VEGF expression, and the involvement of anatomical barriers^60^, they share similar VEGF-driven mechanisms and VEGFR2 signaling, which induces tip cell filopodia formation and guides the growing vessels^61^. Targeted therapies, such as anti-VEGF agents and VEGFR2 tyrosine kinase inhibitors, which have shown efficacy in reducing neovascularization in both models^50, 62^ and have translated into successful treatments for retinopathy of prematurity and neovascular age-related macular degeneration in clinical practice^63, 64^.

This study of EMCN involvement in retinal vasculature development and its impact on pathological angiogenesis models, including OIR and CNV, underscores the important regulatory role of EMCN in angiogenesis. These findings highlight the potential of EMCN as a specific target for anti-angiogenic therapies focused on endothelial cells. These findings pave the way for further exploration of the mechanisms and therapeutic implications of EMCN in the treatment of angiogenesis-related disorders.

